# Exploring the rules of chimeric antigen receptor phenotypic output using combinatorial signaling motif libraries and machine learning

**DOI:** 10.1101/2022.01.04.474985

**Authors:** K.G. Daniels, S. Wang, M.S. Simic, H.K. Bhargava, S. Capponi, Y. Tonai, W. Yu, S. Bianco, W.A. Lim

## Abstract

Chimeric antigen receptor (CAR) costimulatory domains steer the phenotypic output of therapeutic T cells. In most cases these domains are derived from native immune receptors, composed of signaling motif combinations selected by evolution. To explore if non-natural combinations of signaling motifs could drive novel cell fates of interest, we constructed a library of CARs containing ∼2,300 synthetic costimulatory domains, built from combinations of 13 peptide signaling motifs. The library produced CARs driving diverse fate outputs, which were sensitive to motif combinations and configurations. Neural networks trained to decode the combinatorial grammar of CAR signaling motifs allowed extraction of key design rules. For example, the non-native combination of TRAF- and PLCγ1-binding motifs was found to simultaneously enhance cytotoxicity and stemness, a clinically desirable phenotype associated with effective and durable tumor killing. The neural network accurately predicts that addition of PLCγ1-binding motifs improves this phenotype when combined with TRAF-binding motifs, but not when combined with other immune signaling motifs (e.g. PI3K-or Grb2-binding motifs). This work shows how libraries built from the minimal building blocks of signaling, combined with machine learning, can efficiently guide engineering of receptors with desired phenotypes.

**Graphical Abstract:** 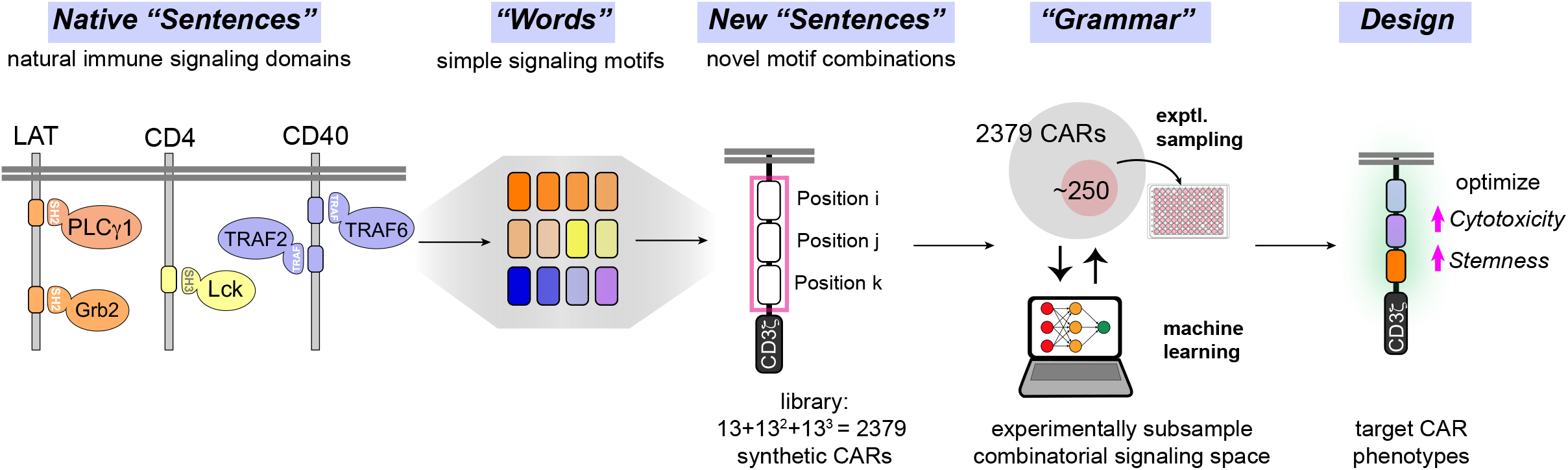

## INTRODUCTION

Chimeric antigen receptors (CARs) have demonstrated the power of synthetic signaling receptors as tools to reprogram immune cells to carry out novel therapeutic functions, such as selective killing of tumor cells (*1*). The antitumor efficacy of CARs is strongly modulated by the signaling domains that they contain. Current clinically approved CARs contain a core TCR signaling domain from CD3ζ (containing ITAM motifs that recruit the kinase ZAP70) (*2-4*), along with a costimulatory signaling domain from either the CD28 (*5, 6*) or 4-1BB (*7*) costimulatory immune receptors (*8-10*). The costimulatory domains are themselves composed of multiple signaling motifs, short peptides that bind to specific downstream signaling proteins, often through modular protein interaction domains (e.g. SH2, SH3, or other domains (*11, 12*)). Such peptide signaling motifs (referred to as linear motifs) are the fundamental building blocks of most signaling receptors. The constellation of signaling proteins recruited by a particular array of signaling motifs is postulated to shape the distinct cellular response. For example, in CARs, the 4-1BB costimulatory domain which contains TRAF binding motifs, leads to increased T cell memory and persistence; the CD28 costimulatory domain, which contains PI3K, Grb2, and Lck binding motifs, is associated with more effective T cell killing, but reduced long-term persistence (*13*). Thus, signaling motifs can be thought of as the “words” that are used to compose the phenotypic “sentences” of signaling domains.

A major goal in synthetic biology is to predictably generate new cell phenotypes by altering receptor composition. For example, in cancer immunotherapy, a general goal is to enhance T cell anti-tumor cytotoxicity as well as maintenance of a stem-like state associated with longer-term T cell persistence. Such a phenotype is associated with effective and durable tumor clearance (higher stemness is correlated with more resistance to T cell exhaustion). In recent work, libraries of costimulatory domains have been screened for improved phenotypes (*14-16*). Such studies, however, have focused on screening intact costimulatory domains from natural immune receptors (i.e. alternative pre-existing “sentences”). We postulate that a more effective way to scan phenotypic space for synthetic receptors, is to create libraries that sample new combinations of signaling motif “words”. Such an approach could, in principle, yield novel phenotypes that extend beyond those that can be generated by native receptor domains alone. Moreover, exploration of a broader range of receptor “motif space” could lead to a more systematic understanding of how different parameters of output are encoded by motif identity, combination and order.

Here we recombine 13 signaling motifs (words) to create a CAR costimulatory domain library with novel motif combinations (new sentences). We show that this library of new signaling “sentences” produces a range of phenotypes, including combination phenotypes not observed with native signaling domains. We then use neural networks to decode the language of signaling motifs, create predictive models, and extract design rules that inform the engineering of CAR signaling domains that increase cytotoxicity and stemness.

## RESULTS

To construct a combinatorial library of CAR signaling domains, we first searched the Eukaryotic Linear Motif Database (ELM)(*17*) and primary literature to curate a collection of 12 peptide motifs from natural signaling proteins known to recruit key downstream signaling proteins involved in T cell activation. The motifs in the library recruit proteins such as PLCγ1, TRAF1/2/3/5, TRAF6, Grb2, GADS, SHP-1, Vav1, PI3K, Lck, and Pellino protein. For example library motif 1 is derived from LAT and contains the core motif YLVV—which binds the N-terminal SH2 domain of PLCγ1 with high specificity(*18*). Motif 6, contains the motif ITYAAV from the protein LAIR1, which binds the phosphate SHP-1 via its SH2 domain (*19*). In addition to the 12 signaling motifs, we included a spacer motif as the 13^th^ component in the library. The combinatorial library was constructed within the context of an anti-CD19 CAR (containing an anti-CD19 extracellular scFv and a CD3 ζ signaling domain). The synthetic costimulatory domains had either one, two, or three signaling motifs. The 13 motifs were randomly inserted in positions i, j, and k to yield 2379 unique motif combinations (Figure 1A-D). To confirm that the library displayed sufficient phenotypic diversity, we first performed low resolution pooled screens, in which we transduced primary human T cells at low MOI and activated the pool with Nalm 6 leukemia cells (CD19+) for 8-9 days. Using FACS-based sequencing enrichment assays, we observed a diverse range of phenotypic outputs for T cell proliferation, central memory formation (CD45RA+/CD62L-T cells), and degranulation (CD107A+ T cells, a proxy for cytotoxic response) (Fig. S1).

**Figure 1.**
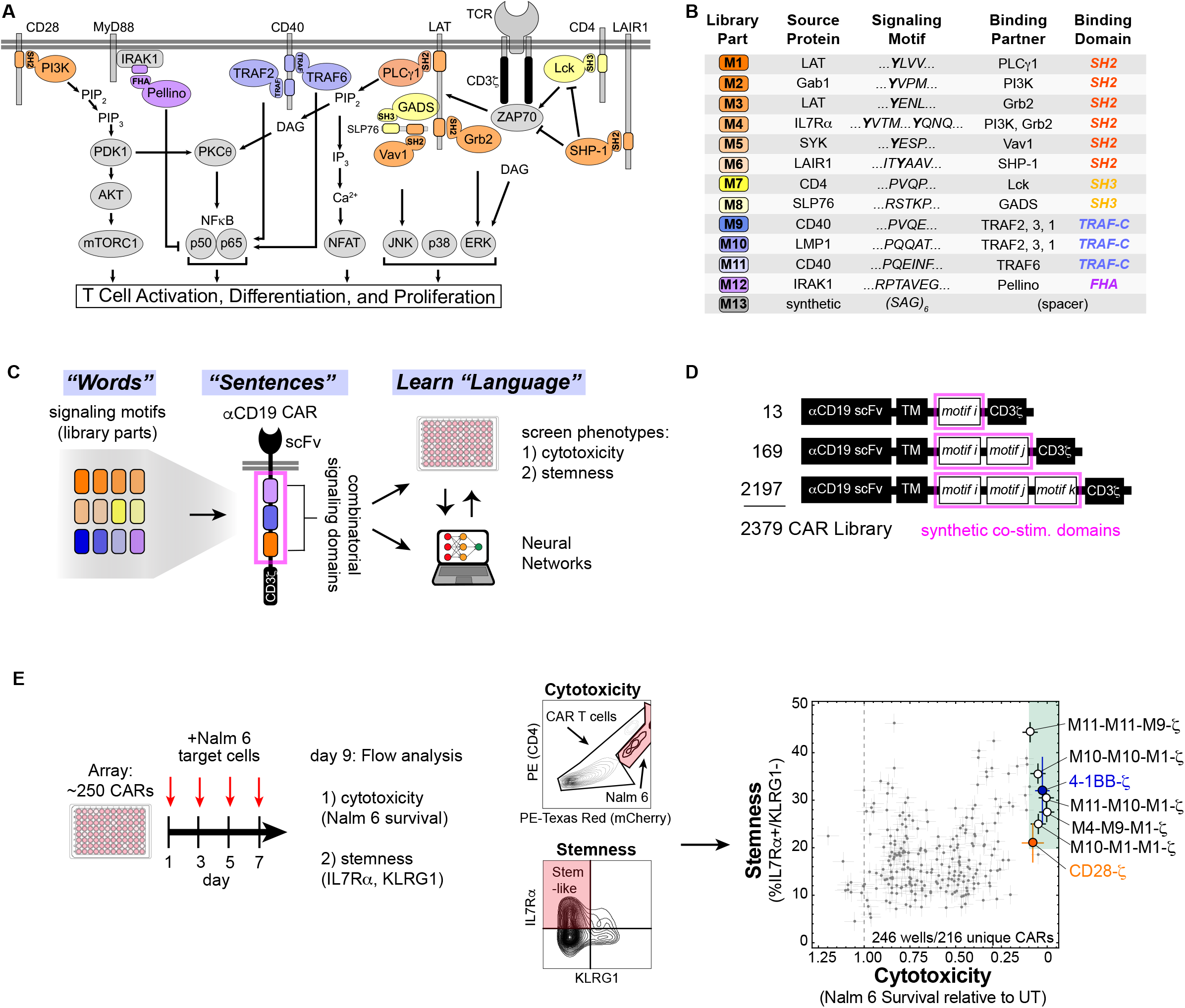
CAR costimulatory domains with novel signaling motif combinations generate diverse cell fates with decoupled cytotoxicity and stemness. **A**, A diverse set of proteins involved in T cell signaling are recruited by signaling motifs in the library parts. **B**, Description of library parts used in combinatorial library. Each part is 16-18 amino acids including the signaling motif(s) and flanking sequence. Phospho-tyrosines are shown in bold. **C**, New combinations of signaling motifs create novel CAR signaling programs that control T cell phenotype. **D**, Schematics of αCD19 CAR with variable signaling domains. **E**, CAR T cells with novel signaling motif combinations produce a broad range of cytotoxicity and stemness. Several combinations produce cytotoxicity and stemness comparable to or exceeding that of CD28 and 4-1BB. Errors for Nalm 6 survival, and stem-like IL7Rα+/KLRG1-population in **E** were estimated by calculating the average s.e.m. for 7 CAR constructs with internal duplicates in the array.

To then screen the library at higher resolution, we transformed bacteria with library plasmid stocks and randomly picked colonies to select a subset of over 200 novel CARs from the combinatorial library to characterize in an arrayed screen (Fig. 1F). An arrayed screen was important because immune paracrine signaling could confound analysis of mixed CAR T cell populations and allow the emergence of “cheater” individuals. We activated the CAR T cells in the arrayed screen by co-culturing with Nalm 6 cells for 8-9 days. Four pulses of Nalm 6 cells were used to mimic longer term stimulation that can exacerbate T cell exhaustion. At the end of the co-culture, we performed flow cytometry to assess cytotoxicity (Nalm 6 cell killing), and maintenance of T cell populations with markers of central memory/naïve state (CD45RA and CD62L) and stemness (IL7Rα and KLRG1) (*20, 21*).

The CARs in the arrayed screen displayed a range of cytotoxicity as well as stemness. The total naïve and central memory population was positively correlated with cytotoxicity (Fig. S2). However, cytotoxicity and stemness were uncoupled. This observation underscores the ability of novel costimulatory domains to drive CAR T cells to varied cell fates with unique combinations of phenotypes. Several novel costimulatory domains produced cytotoxicity and stemness comparable to 4-1BB. Many of these contained motifs that recruit both TRAFs (motif 9, motif 10, motif 11) and PLCγ1 (motif 1). For example M10-M10-M1-ζ, M10-M1-M1-ζ, M11-M10-M1-ζ, and M4-M9-M1-ζ all showed robust cytotoxicity and stemness.

The diverse cytotoxicity and stemness profiles observed in our arrayed screen suggest a complex relationship between signaling motif combinations and arrangement, and resulting T cell phenotypes. We sought to leverage the combinatorial nature of the costimulatory domain library by using machine learning to decode the “language” of signaling motifs that relates motif combinations to cytotoxicity and stemness outputs. We separated the arrayed screen data into a training sets (221 examples) and a test set (25 examples). We then used these data sets to train several machine learning algorithms to predict cytotoxicity and stemness based on costimulatory domain identity and arrangement (Figs. S3, 2A). Neural networks (Figure 2B) were best able to recapitulate the measured phenotypic in the training data (Figure 2C) and to effectively predict the phenotypes in the test set (Figure 2D). For both cytotoxicity and stemness training and test sets, the neural network was able to capture much of the relationship between signaling motif composition and phenotype, with R^2^ values of approximately 0.7-0.9.

**Figure 2.**
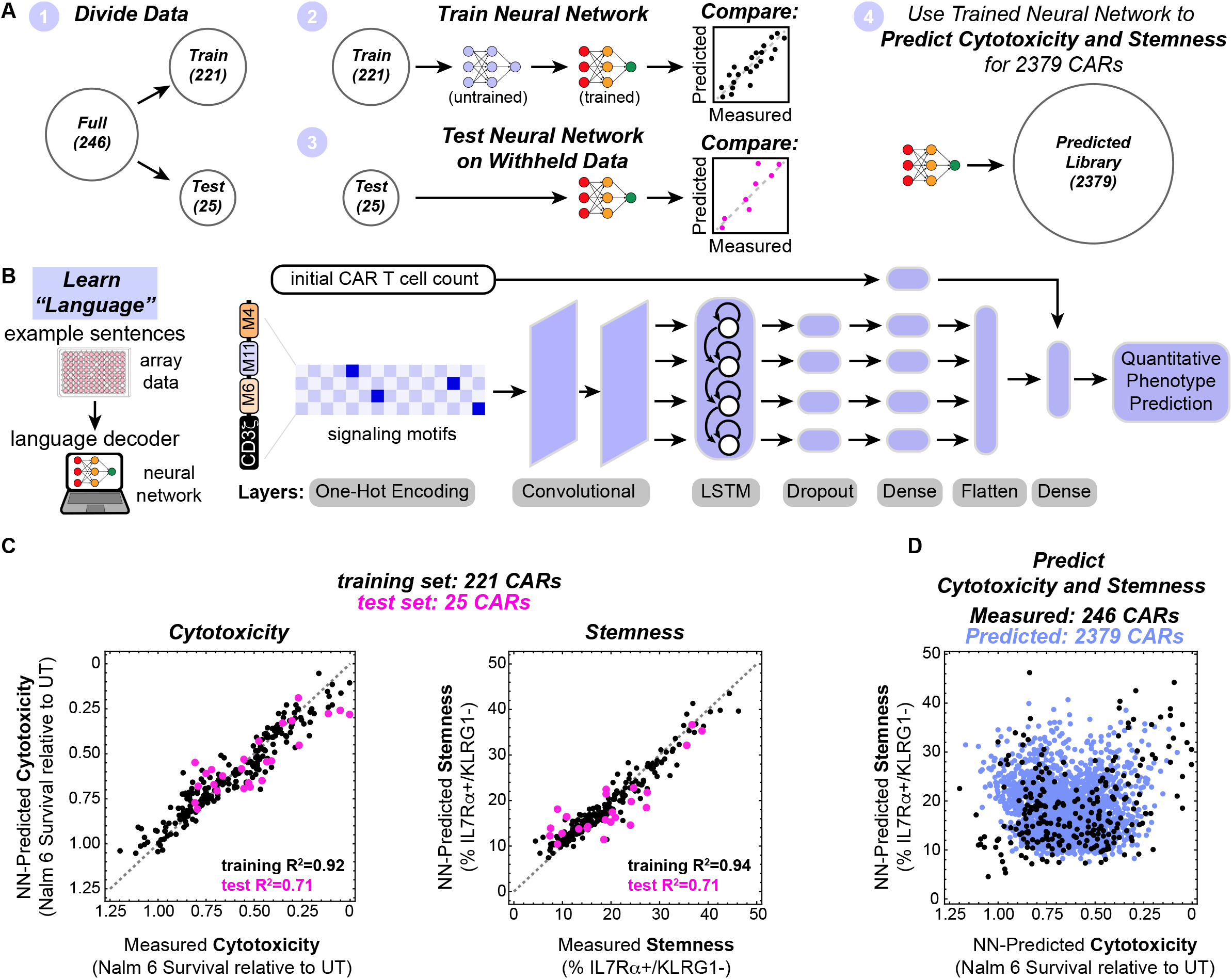
Neural networks decode the combinatorial language of signaling motifs to predict cytotoxicity and stemness of novel motif combinations. **A**, Array data were subdivided in datasets to train and test neural networks that were subsequently used to predict the cytotoxicity and stemness of 2379 CARs. **B**, Schematic of neural network used to predict CAR T cell phenotype. **C**, Neural networks trained on array data predict the cytotoxicity and stemness of CARs in the training sets (black) and the withheld test sets (pink). **D**, Trained neural networks were used to predict the cytotoxicity and stemness of 2379 CARs containing 1-3 variable signaling motifs. Predictions represent the mean for n=10 neural networks with different hyperparameters.

The trained neural networks then allowed us to predict the CAR T cell cytotoxicity and stemness that would result from each of the 2379 motif combinations in the 1-3 part combinatorial library (Figure 2D), including those that were not part of the smaller arrayed screen. These simulated 2379 CARs sample the entire combinatorial space of the library, providing a dataset from which we could extract design rules. Below we describe three types of analysis: 1) the overall contribution of each individual motif to a particular phenotype (without regard to combinatorial context); 2) identification of pairwise motif combinations that promote particular phenotypes, and 3) positional dependence of motifs.

To assess the overall contribution of individual motifs, we ranked all the CARs in our library by neural network predicted cytotoxicity and stemness and then assessed whether motifs were enriched in the strong or weak ends of the phenotypic distribution (Figure 3A, S4). If a motif is generally activating for a phenotype, then it is expected to be more common in highly ranked CARs; if a motif is inhibitory it is expected to be more common in poorly ranked CARs. Although the effects observed in this distribution analysis depend on other motifs in the CAR and the position of the motif in question, the distributions are informative of the overall effect each motif has in the context of the library. An analogous distribution analysis was also performed on the pooled screening proliferation data (Figure S5).

**Figure 3.**
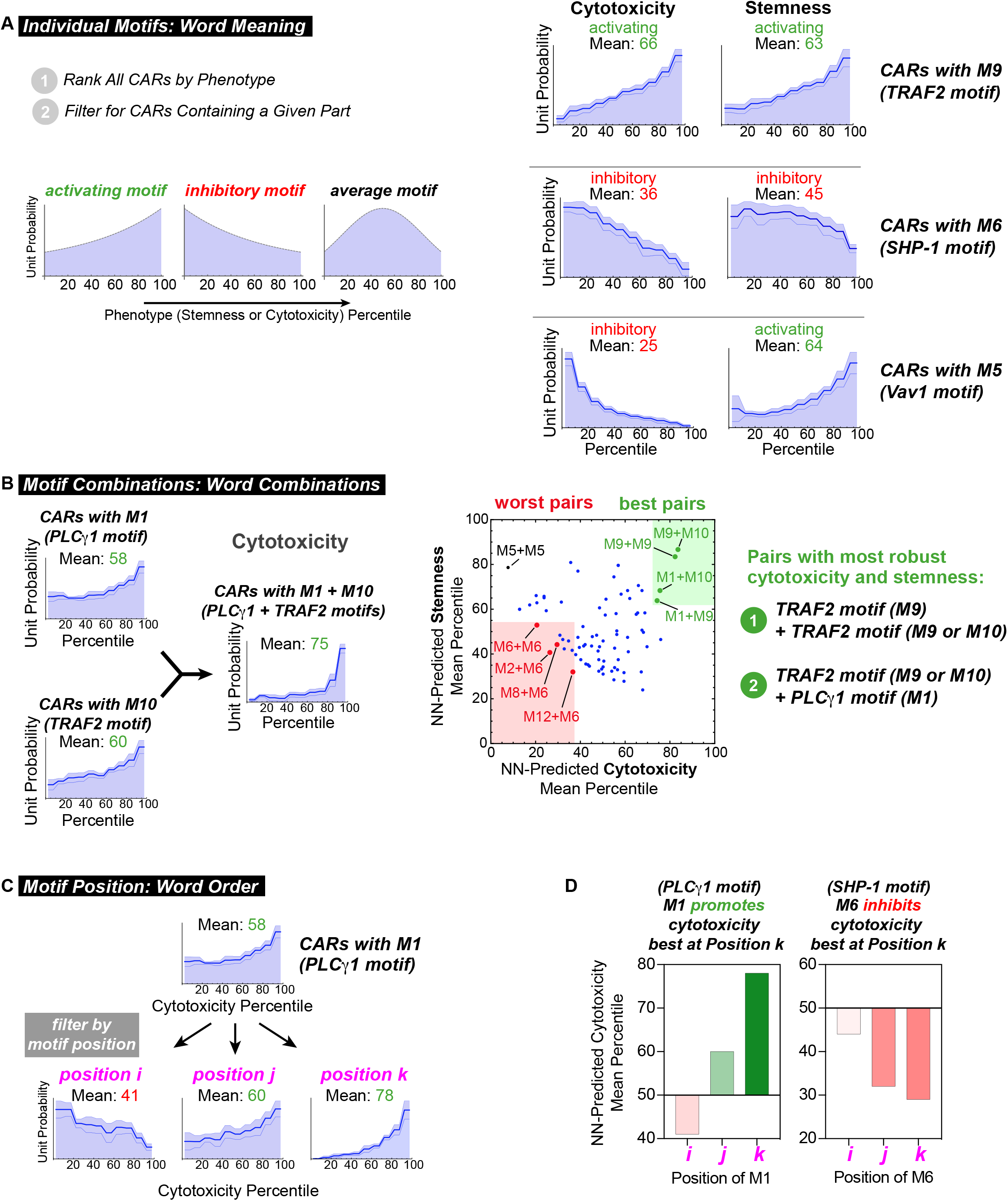
Distribution analysis quantifies elements of linear motif language to extract design parameters for signaling domains. **A**, The distribution of library parts throughout CARs in the ranked library reflects effects of signaling motifs on phenotype. Activating motifs are found in CARs with higher rank and inhibitory motifs are found in CARs with lower rank. The three lines within the distributions represent mean predictions ± s.e.m. calculated from n=10 neural networks. **B**, CARs containing pairs of motifs that recruit TRAFs (P9 and P10) or PLCγ1 (P1) promote robust cytotoxicity and stemness. **C**, The effects of signaling motifs on phenotype are position-dependent. **D**, P1 is predicted to promote cytotoxicity best at position k, while P6 is predicted to inhibit cytotoxicity best at position k.

This distribution analysis highlighted several potent motifs know to play activating and inhibitory roles. For example motif M9 is the PQVE motif (from CD40) that binds TRAF2, and is associated with T cell activation and function(*22, 23*). Accordingly, M9 is enriched in CARs with high cytotoxicity (mean 66 percentile) and high stemness (mean 63 percentile), indicating that overall it promotes both of these phenotypes. In a contrasting example, M6 (from LAIR1) recruits the phosphatase SHP-1, a potent inhibitor of T cell activation. Accordingly, M6 is enriched in CARs with low cytotoxicity (mean 36 percentile) and low stemness (mean 45 percentile), indicative of inhibition of both phenotypes. Interestingly, some motifs can activate one phenotype and inhibit another: M5, which binds Vav1, is unrepresented in CARs with high cytotoxicity (mean 25 percentile), but overrepresented in CARs with high stemness (mean 64 percentile). These results suggest the unexpected finding that Vav1 signaling promotes stemness, while inhibiting killing. The quantified effect of all individual motifs on cytotoxicity and stemness are shown in the heatmap in Figure S4. The TRAF binding motifs (M9 and M10) are among the best at promoting both cytotoxicity and stemness.

We anticipated that phenotypes would be highly dependent on motif combinations, as different downstream signaling pathways could be either complementary or redundant/competing. To examine motif pairs that favored particular phenotypes, we examined the occurrence of each possible pair (without regard to order) in the ranked distribution. Several specific motif pairs appear to promote both robust cytotoxicity and stemness when they occur in combination within a costimulatory domain. For example, M1 (PLCγ1) and M10 (TRAF) are each activating with respect to cytotoxicity (means: 58 and 60), but the P1+P10 motif pair is even more strongly activating (mean: 75 percentile). The predicted mean cytotoxicity and stemness percentiles for all 144 pairs of motifs M1-M12 is shown in Figure 3B. The motif pairs M1+M10, M1+M9, M9+M9, and M9+M10 are best at promoting cytotoxicity and stemness. These pairs all suggest that TRAF-binding motifs (M9 and M10) work well in tandem, as well as in combination with the motif that recruits PLCγ1 (M1) whose signaling is known to activate NFκB. As examined below, we hypothesize that these pathways are likely to serve complementary roles in these phenotypes. A number of motif pairs strongly inhibit cytotoxicity and stemness. Not surprisingly, all four motif pairs with the lowest cytotoxicity and stemness contain M6 which binds the inhibitory phosphatase SHP-1. Similarly, the tandem occurrence of M5 (Vav1 motif) leads to the strongest combination of stemness with low cytotoxicity.

Finally, we also find that phenotype can be highly dependent on the position of a motif within the costimulatory domain (Figure S4B). For example, M1 (PLCγ1) shows strong cytotoxicity when in positions k or j, and weak cytotoxicity in position i (Fig. 3C). M9 (TRAF) and M10 (TRAF) show optimal cytotoxicity and stemness when in positions i and j. This is consistent with the experimental observation that TRAF-binding parts M9 and M10 followed by M1 (in N-to C-terminal order) promote the most robust cytotoxicity and stemness (M1 followed by M9 or M10 does not (Fig. S2C). These results suggests that shuffling motif position is an approach for calibrating phenotype.

The above distribution analysis quantifies elements of motif language, capturing the effects of motifs (word meaning), motif pairs (word combinations), and motif position (word order) on phenotype. The analysis also yields design rules that can inform combinations and arrangements of motifs capable of producing a desired cell fate. For example, a clearly emergent hypothesis is that a synthetic costimulatory domain which contains one or more TRAF binding motifs (M9 or M10) followed by a PLCγ1 (M1) motif could be highly effective at promoting both cytotoxicity and stemness (Fig. 4A). While tandem TRAF binding motifs occur in the naturally evolved 4-1BB receptor (*24*) (Fig. S6A), the combination of TRAF and PLCγ1 motifs are not found in natural characterized immune receptors. Thus, we wanted to test if adding PLCγ1 (M1) motifs to 4-1BB-like domains could improve CAR phenotype. Moreover, we also wanted to determine if adding M1 might be a general strategy to improve the efficacy of other costimulatory domains, such as CD28.

**Figure 4.**
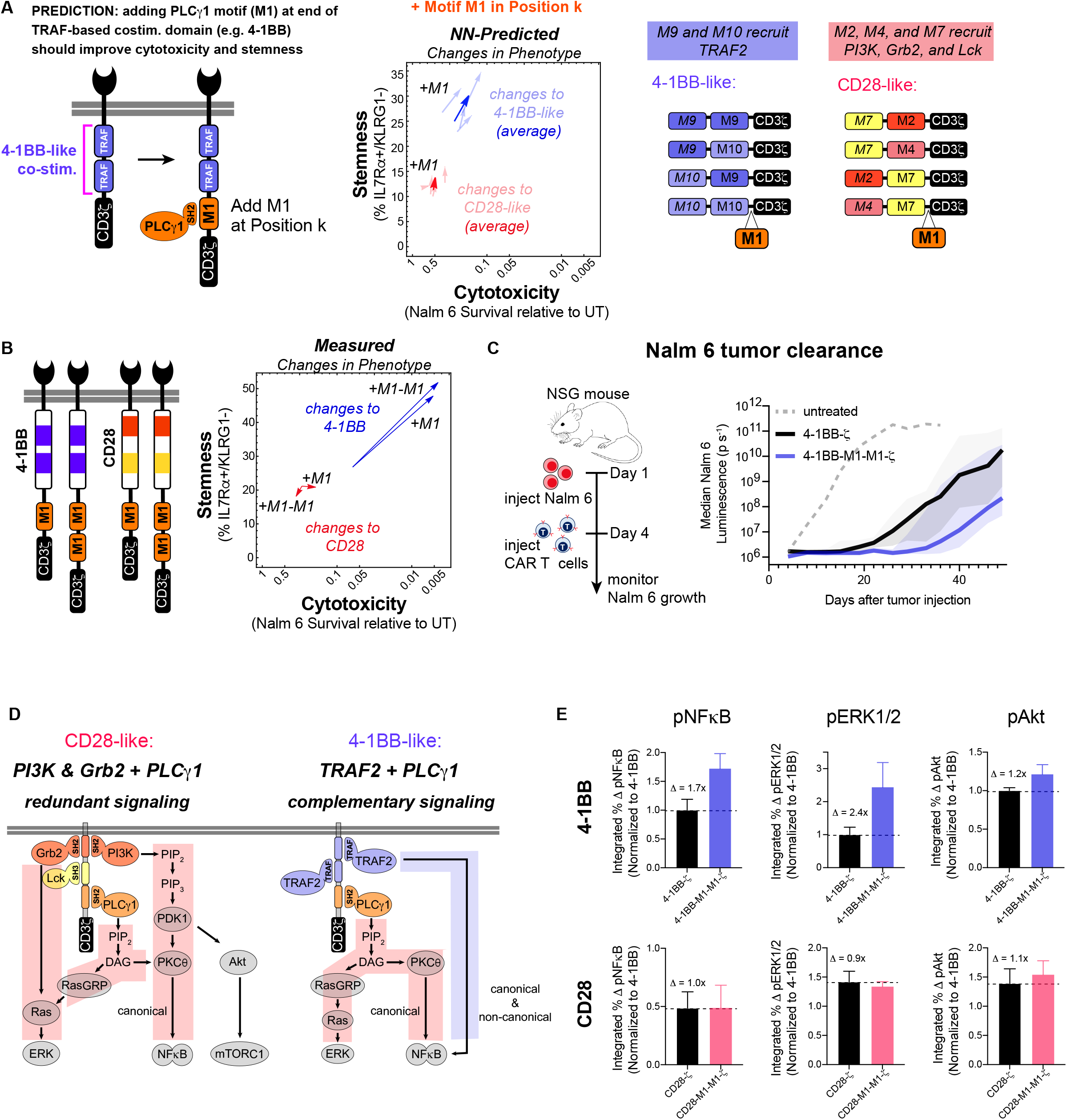
Neural networks accurately predict that PLCγ1 binding motifs improve the cytotoxicity and stemness of 4-1BB-ζ but not CD28-ζ. **A**, Library parts that share consensus signaling motifs with 4-1BB and CD28 costimulatory domains were used to predict the effect of adding at P1 to 4-1BB and CD28. **B**, Addition of 1 or 2 copies of P1 improved i*n vitro* cytotoxicity and stemness of 4-1BB-ζ but not CD28-ζ. Cytotoxicity and stemness were assessed after 4 challenges with Nalm 6 cells. Data are mean ± s.e.m. of *n* = 3-5 replicates. **C**, NSG mice were injected intravenously with 0.5 × 10^6^ Nalm 6 cells, and then injected intravenously with 3 × 10^6^ CAR^+^ T cells on day 4. CAR T cells with 4-1BB-P1-P1-ζ showed improved early tumor control relative to 4-1BB-ζ. Traces in **C** are median luminescence ± 95% confidence interval. **D**, Costimulatory PLCγ1 signaling is redundant to signaling provided by PI3K and Grb2, but complementary to TRAF signaling. Addition of P1 to 4-1BB-ζ induced modest changes in Akt phosphorylation—which is not downstream of PLCγ1 signaling—relative to the changes in ERK1/2 and NFκB phosphorylation—which are downstream of PLCγ1 signaling. Data for **D** are mean and standard deviation of *n* = 3 replicates.

To explore this hypotheses, we first examined the neural network-predicted library to predict the effects of adding the M1 motif to CD28-like and 4-1BB-like synthetic costimulatory domains (library members whose signaling motifs shared the overall configuration of natural signaling motifs in CD28 and 4-1BB) (Fig. 4A). The 4-1BB-like costimulatory domains are predicted by the neural network model to show increased cytotoxicity and stemness, consistent with experimental observations. In contrast, addition of M1 motifs to CD28-like costimulatory domains are not predicted to enhance cytotoxicity or stemness.

Next, to experimentally test the hypothesis, we synthesized derivatives of the 4-1BB and CD28 costimulatory domains with 1 or 2 copies of the M1 motif added to the C-terminus, and tested these costimulatory domains for their ability to kill Nalm 6 and maintain stemness (Fig. 4B). Consistent with predictions, 4-1BB showed significantly enhanced cytotoxicity and stemness upon addition of M1, while CD28 showed almost no change. Significantly, in addition to predicted in vitro changes, the 4-1BB-M1-M1-ζ CAR construct showed improved efficacy against a Nalm 6 tumor NSG mouse model (Fig. 4C, Fig. S6D). Relative to standard 4-1BB CAR T cells, the 4-1BB-P1-P1-ζ CAR T cells were able to delay the growth of Nalm 6 for an additional two weeks, validating the predictions from the library/neural network model.

Why might a PLCγ1 motif improve T cell phenotype when in combination with the 4-1BB domain (TRAF motifs) but not in the context of the CD28 domain (PI3K, Grb2, Lck motifs)? PLCγ1 catalyzes the production of DAG from PIP_2_, which activates RasGRP and PKCθ, subsequently activating ERK1/2 and NFκB. This signaling is similar and possibly redundant to that of PI3K and Grb2, which also activate RasGRP and PKCθ. TRAF signaling, however, does not activate RasGRP or PKCθ, such that PLCγ1 and TRAF signaling are likely to be more complementary (Fig. 4D). We experimentally characterized the 4-1BB-M1-M1-ζ CAR construct (compared to standard 4-1BB-ζ CAR) by measuring the kinetics of Akt, ERK1/2, and NFκB phosphorylation upon Nalm 6 stimulation (Fig. 4E, Fig. S6E). The addition of the M1 motifs increase phosphorylation of ERK1/2 (1.7-fold) and NFκB (2.4-fold), both of which are downstream of PLCγ1 activity. Phosphorylation of Akt, which is not downstream of PLCγ1 activity, shows only a 1.2-fold increase. The observed increase in NFκB and ERK1/2 activation support the hypothesis that PLCγ1 signaling is complementary to TRAF signaling and is consistent with previous reports that NFκB activation is important for the maintenance of CD8+ T cell memory(*25*). In contrast, no significant increase in NFκB and ERK1/2 activation is observed for a CAR in which the PLCγ1 motif is appended to the CD28 costimulatory domain.

## CONCLUSIONS

In conclusion, we find that signaling motif libraries and machine learning can be combined to elucidate rules of CAR costimulatory signaling and to guide the design of novel, non-natural costimulatory domains with improved phenotypes, both in vitro and in vivo. Costimulatory signaling modulates the outcome of CAR T cell activation, making costimulatory domains attractive engineering targets for customizing or improving cell therapies. Thus far, costimulatory domain engineering has mostly been limited to the addition of intact natural domains such as 4-1BB, CD28, or the IL2Rβ chain, effectively using naturally occurring signaling sentences (motif combinations). Here we used motifs from receptors as words to generate thousands of novel signaling sentences that drive T cells to distinct cell fates, potentially yielding more diverse and nuanced phenotypic meaning. Augmenting experimental analysis of a subset of receptors with neural network analysis allows us to explore a much larger region of this combinatorial motif space. In particular, we identified that the non-natural combination of TRAF- and PLCγ1-binding motifs that may be useful in CAR T cell therapies. By performing an arrayed screen of several hundred receptors and using machine learning, we identified basic elements of signaling motif language and extracted design rules that relate motif combinations to cell fate. This represents a step toward forward engineering receptors with desired properties. Exploration of larger libraries will benefit greatly from machine learning due to the size and complexity of the combinatorial space. Future machine learning-augmented screens of this type could be used engineer many other classes of receptors for biological research and cell therapy applications involving cellular processes controlled by combinations of signaling motifs.

## METHODS

### Viral vector construction

Codon-optimized DNAs encoding the variable library parts were codon optimized for expression in human cells using ThermoFisher GeneArt’s website tool and synthesized by ThermoFisher GeneArt. A pHR lentiviral vector containing an SFFV promoter followed by DNA encoding the aCD19 scFv and the CD8a hinge and transmembrane domain was BamHI restriction digested. DNA encoding a BamHI cut site followed by CD3z-P2A-EGFP was subcloned into the digested pHR-aCD19-(BamHI)-CD3z-P2A-EGFP vector to create library backbone. DNA encoding library parts, as well as DNA encoding 4-1BB, CD28, and ICOS, was subcloned into the library backbone and the cloning product was again BamHI digested. This was repeated to create vectors with 3 variable library parts. All constructs were built via in-fusion cloning (Clontech #ST0345) and sequence verified before use. Amino acid sequences for library parts were as follows. P1: DYHNPGYLVVLPDSTP, P2: EELDENYVPMNPNSPP, P3: EEGAPDYENLQELNHP, P4: LGSNQEEAYVTMSSFYQNQ, P5: LPMDTEVYESPFADPEEIR, P6: KPMAESITYAAVARHSAG, P7: LPTWSTPVQPMALIVLG, P8: PAPSIDRSTKPPLDRSL, P9: GSNTAAPVQETLHGCQ, P10: DDSLPHPQQATDDSGHES, P11: KAPHAKQEPQEINFPDDLP, P12: GSGPGSRPTAVEGLALGSS, P13: SAGSAGSAGSAGSAGSAG.

### Cell lines

Nalm 6 cell lines were originally obtained from ATCC (CRL-3273) and were stably transduced with mCherry and firefly luciferase. Nalm 6 cell lines were cultured in RPMI-1640 + GlutaMAX (Gibco #72400-047) supplemented with 10% FBS (UCSF Cell Culture Facility).

#### Primary human T cell isolation and culture

Primary CD4+ and CD8+ T cells were isolated from blood of anonymous donors by negative selection using the Human CD4+ T cell isolation kit and Human CD8+ T cell isolation kit (STEMCELL Technologies #17952 and #17953). T cells were cryopreserved in RPMI1640 (UCSF cell culture core) with 20% human AB serum (Valley Biomedical, #HP1022HI) and 10% DMSO. Upon thawing, T cells were cultured in human T cell medium (HTCM) consisting of X-VIVO 15 (Lonza #04-418Q), 5% Human AB serum, 1 mM 2-mercaptoethanol (Gibco #21985-023), and 10 mM neutralized N-acetyl L-Cysteine (Sigma-Aldrich #A9165) supplemented with 30 units/mL IL-2 (NCI BRB Preclinical Repository). Before co-culture with Nalm 6, T cells were transferred to HTCM without IL-2.

### Lentiviral Transduction and Sorting of Human T Cells

Pantropic vesicular stomatitis virus G pseudotyped lentivirus was produced via transfection of LentiX 293T cells (Clontech #11131D) with a pHR’SIN:CSW transgene expression vector and the viral packaging plasmids pCMVdR8.91 and pMD2.G using FuGENE HD (Promega, #E2312). Primary T cells were thawed the same day and after 24 h in culture, were stimulated with Dynabeads Human T-Activator CD3/CD28 (Life Technologies #11131D) at 25 μL per 1 × 10^6^ T cells. At 48 h (day 2), viral supernatant was harvested via centrifugation at 500 G for 5 min, and the primary T cells were exposed to the virus for 24 h in a 6-well plate (pooled screens) or in 96-well plates (arrayed screens). Dynabeads were removed at day 5 post-T cell stimulation. For pooled screens, GFP+ T cells were sorted on day 6 post-T cell stimulation with a FACSAria II. Assays were performed 10 days after removal of Dynabeads.

### Arrayed Screening

CARs were constructed as described above in Viral Vector Construction. An additional pooled CAR sub-library was constructed with enriched concentration of DNA corresponding to P1, P4, P7, P9, and P10 on the basis of their high proliferation, degranulation, and memory formation in the pooled screening assay. Pooled CAR library DNA was used to transform 5-alpha F’ I^q^ competent E. coli cells (New England BioLabs C2992H), which were then plated on LB/Carbenicillin. At 24 hours 384 colonies (288 from the unbiased library, and 96 from the high-performance sub-library) were picked and miniprepped, added to 96-well plates and sequence verified. Wells with failed sequencing results or unidentifiable sequences were removed from plates and the well contents were replaced with duplicates of nearby wells, TE buffer (for empty well controls), or standard costimulatory domain (4-1BB, CD28) controls. CARs containing 4-1BB, CD28, and ICOS costimulatory domains and P1-P13 were left in place, but excluded from the analysis.

Primary human T cells transduced with CAR library constructs were mixed with Nalm 6 to reach 1×10^6^ T cells per mL and 2×10^6^ per mL Nalm 6 and centrifuged at 300g for 2 min. For day 3, 5, and 7 challenges with Nalm 6, 80 μL of co-cultured T cells and Nalm 6 were added to 120 μL of Nalm 6 at 2×10^6^ per mL and centrifuged at 300g for 2 min.

For analysis of cell surface receptor expression, samples were centrifuged at 500 × g for 5 min and resuspended in a 50 μL volume with the appropriate antibodies diluted 1:50 in calcium-free magnesium-free PBS with 5% FBS and 5mM EDTA. After a 30-min incubation at room temperature, samples were washed twice calcium-free magnesium-free PBS with 5% FBS and 5mM EDTA. (FBS; UCSF Cell Culture Facility). Samples were analyzed for protein expression on a BD LSRII. Antibodies are as follows: APC Mouse anti-human KLRG1 clone SA231A2 (BioLegend #367716), BV421 Mouse anti-human IL7Rα clone HIL-7R-M21 (BD Biosciences #562436), BV786 Mouse anti-human CD62L clone SK11 (BD Biosciences #565311), AF700 Mouse anti-human CD45RA clone HI100 (BD Biosciences #560673), PE Mouse anti-human CD4 clone RPA-T4 (BD Biosciences #555347).

### Pooled Screening

Primary human T cells transduced with pooled virus for the CAR library were mixed with Nalm 6 to reach 1×10^6^ T cells per mL and 2×10^6^ per mL Nalm 6. For day 3, 5, and 7 challenges with Nalm 6, co-cultured T cells and Nalm 6 were centrifuged at 400g for 4 min and resuspended at 1×10^6^ per mL in 1/3 current HTCM and 2/3 fresh HTCM. Additional Nalm 6 were added at 2×10^6^ per mL.

For extracellular staining, samples were centrifuged at 500g for 5 minutes and resuspended in FACS buffer with 1:50 PE anti-human CD4 antibody, 1:50 BV421 mouse anti-human CD45RA antibody, and 1:50 AF647 mouse anti-human CCR7 antibody. After a 30-min incubation at room temperature, samples were washed twice, and resuspended in FACS buffer. Samples were sorted on a BD FACS AriaII.

For analysis of degranulation on Day 9 Nalm 6 challenge, samples of 1×10^6^ per mL pooled CAR T cells with 2×10^6^ per mL Nalm 6 were centrifuged at 300 × g for 2 min in 96-well flat-bottom plates and incubated in HTCM with 1x Brefeldin-A/GolgiPlug (BDBiosciences #555029), 1x Monensin/GolgiStop (BD Biosciences #554724), and 1:50 APC anti-human CD107A antibody at 37C and 5% CO_2_ for 5 hours. At 5 hours, samples were centrifuged at 300g for 2 minutes and supernatant was removed. Samples were resuspended in FACS buffer with 1x GolgiPlug, 1x GolgiStop, and 1:50 PE anti-human CD4 antibody. After a 30-min incubation at room temperature, samples were washed twice, and resuspended in FACS buffer. Samples were sorted on a BD FACS AriaII. Antibodies were as follows: PE Mouse anti-human CD4 clone RPA-T4 (BD Biosciences #555347), BV421 Mouse anti-human CD45RA clone HI100 (BD Biosciences #562885), AF647 Mouse anti-human CCR7 clone 150503 (BD Biosciences # 560816), APC Mouse anti-human CD107A clone H4A3 (Biosciences #641581).

Genomic DNA was extracted from sorted T cells using the Macherey-Nagel NucleoSpin Tissue XS kit (Takara #740901.250). DNA encoding the CAR costimulatory domain was amplified from the extracted genomic DNA using the forward primer 5’-TCGTCGGCAGCGTCAGATGTGTATAAGAGACAGNNNNNACTGGTTATCACCCTTTA CTGC-3’ (Integrated DNA Technologies) and reverse primer 5’-GTCTCGTGGGCTCGGAGATGTGTATAAGAGACAGNNNNNCTTGTAGGCGGGAGC AT-3’ (Integrated DNA Technologies). Indexes were added to the amplified DNA using i5 and i7 primers from the Nextera XT Index Kit (Illumina # FC-131-1002). Indexed samples were loaded into a MiSeq Reagent Kit v3 600-cycle (Illumina # MS-102-3003) cartridge sequenced on a MiSeq (Illumina). Reads for each CAR costimulatory domain construct were counted using software provided by Ian Webster at Zenysis Technologies.

### Assessment of Akt, ERK1/2, and NFκB phosphorylation

For intracellular phospho-signaling analysis, T cells and Nalm 6 cells were plated at 1:2 ratio in HTCM and centrifuged at 400g for 3 minutes. Cells were plated and centrifuged at t= 0, 30, 45, 50, 55, and 59 min to obtain timepoints for approximately 60, 30, 15, 10, 5, and 1 minutes of T cell:Nalm 6 engagement. After the final sample was centrifuged, all samples were mixed 1:1 with prewarmed CytoFix Fixation Buffer (BDBiosciences #554655) and incubated at room temperature for 15 minutes. Samples were washed twice with FACS buffer, vigorously vortexed, and permeabilized with BD Phosflow Perm Buffer II (BDBiosciences #558050) by incubating overnight at -20C. Permeabilized samples were washed twice with FACS buffer and resuspended in 50 μL of FACS buffer with 1:50 anti-human phospho-NFκB antibody, 1:50 anti-human phospho-Akt antibody, and 1:50 anti-human phospho-ERK1/2 antibody. After a 60-minute incubation, samples were washed twice with FACS buffer and analyzed by flow cytometry on a BD LSRII. Antibodies are as follows: BV421 Mouse anti-human NFκB pS529 clone K10-895.12.50 (BD Biosciences #565446), PerCP-Cy5.5 Mouse anti-human ERK1/2 pT202/pY204 clone 20A (BD Biosciences #560115), PE-CF594 Mouse anti-human Akt pS473 clone M89-61 (BD Biosciences #562465), AF647 Mouse anti-human Akt pS473 clone M89-61 (BD Biosciences #560343).

Flow cytometry data were analyzed in FlowJo (BD) software to calculate mean fluorescence intensity (MFI) for pErk1/2, pNFκB, and pAkt channels. MFI kinetic traces were interpolated and integrated in *Mathematica* (Wolfram) to calculate the total change over 60 minutes in MFI for CAR T cell samples relative to the change in MFI of the untransduced control. Integrated changes in MFI were normalized to 4-1BB measurements to standardize experiments performed on different days.

### Mice

All mouse experimental procedures were conducted according to Institutional Animal Care and Use Committee (IACUC)–approved protocols. Female immunocompromised NOD-SCID-*Il2rg*^™/™^ (NSG) mice were obtained from UCSF breeding core. On Day 1, mice were inoculated with 0.5 × 10^6^ Nalm 6 leukemia via tail vein injection. On Day 4, mice were injected with 3×10^6^ T cells via tail vein injection. Leukemia progression was measured by bioluminescent imaging using the IVIS 100 (Xenogen) preclinical imaging system. Images were acquired 15 minutes following intraperitoneal (i.p.) injection with 150 mg/kg of D-luciferin (Gold Technology #LUCK-100). Display and adjustment of bioluminescence intensities was performed using the Living image 4.5.4 software (Perkin Elmer). Mice were humanely euthanized when IACUC-approved endpoint (hunching, neurological impairments such as circling, ataxia, paralysis, limping, head tilt, balance problems, seizures, tumor volume burden) was reached (10 mice per group).

## Machine Learning

### Data Preparation

For the arrayed data, in addition to the positional information of the combinational motifs, the initial CAR T cell number is a variable which affects the experimental output. Both are inputs of the machine learning algorithms. We randomly split D2 and use 90% for training and 10% for test, (we repeated this splitting process until duplicate motif combinations were found either exclusively in the training sets or exclusively in the test sets) ensuring all the motif combinations in the test data are different from those in the training.

We used one-hot encoding to input motif combinations into the machine learning algorithms. Each motif position was described by a vector of fifteen 0s, and one 0 in each vector was replaced with a 1 corresponding to the absence of a motif (replace the first 0 with 1), the presence of a motif (replace the 0 equal to the part number + 1 with 1), or the presence of CD3z (replace the 15th 0 with 1). We allowed up to 5 motif positions, as well as CD3z, for a total of 6 vectors. This allows for inclusion of a small number of CARs that contained more than 3 motifs, and allows flexible inclusion of additional data for CARs with more than 3 motifs.

### Machine learning Framework

In this work, we used a Convolutional Neural Networks (CNN), followed by a Long Short-Term Memory (LSTM) network together with fully connected layers. The code is implemented in either *Mathematica* (Wolfram) or Python 3.7.8 and TensorFlow v2.4.1, both of which produce nearly identical results. The Mathematica analysis is described below and more detailed analysis is described in the forthcoming companion paper. The neural network uses the AΧB matrices as inputs and outputs one value corresponding to one of the phenotypes (cytotoxicity and stemness). Between the input and output layers, we have 2 convolutional layers, 1 LSTM layer, 1 dropout layer, and several fully connected layers. The convolutional layers detect spatial correlations in input data and the LSTM layer learns the long-term dependencies of the sequence data. We used dropout regularization to prevent over-fitting. The dropout layer connects to fully connected layers which are then flattened and catenated with the cell number input and connect to a dense layer. We used linear activation function to connect this dense layer and the final output layer. For training in Mathematica, we used mean squared error loss and ADAM optimization algorithm with automatic learning rate, and training over 200 iterations.

We also compared our methods with other widely used machine learning regression methods, such as k-nearest neighbor regression, linear regression, nearest neighbors, random forest regression, and gradient boosted regression. Our method has the best performance and predictive power compared to other methods. Details of the analysis can be found in the forthcoming companion paper.

### Selection of neural network Hyperparameters

We tuned the hyperparameters for layers in the neural networks to find optimal hyperparameters for the cytotoxicity and stemness datasets. The tuned hyperparameters include convolutional layers filters (10, 20, 50), kernel size (2, 3, 4, 5); LSTM layer units (2, 4, 8), dropout layer dropout rate (0, 0.1, 0.2), and fully connected layer units (6, 14, 64).

Hyperparameters were tuned as follows: We performed a grid search of hyperparameters and scored each parameter set by 10-fold cross validation of the training set. The best-performing 10 hyperparameter sets for each dataset (cytotoxicity or stemness) were selected using the K-fold averaging cross validation (ACV) method and used to train 10 neural networks whose outputs were then averaged(*26*). The trained neural networks were used to simulate the cytotoxicity and stemness for the 2379 combinations of 1, 2, or 3 variable motifs at a fixed initial cell count of 2000 cells (corresponding to 2000 CAR T cells in 40 μL of flow cytometry sample).

Hyperparameters for final neural networks are available in supplementary tables.

### Ensemble Method

Due to the stochastic nature of network initialization and dropout, as well as the availability of a limited training set, every neural network is unique in terms of the parameterization of the network connections(*27, 28*). To mitigate the potential impact of this issue, we implemented an ensemble decision method to obtain consensus prediction from ten identical neural networks. Details can be found in the forthcoming companion paper.

### Distribution analysis

Distribution analysis was performed in *Mathematica* (Wolfram). CARs were sorted by proliferation (lowest enrichment to highest enrichment), cytotoxicity (highest Nalm 6 survival to lowest Nalm 6 survival), or stemness (lowest %IL7Rα+/KLRG1-to highest %IL7Rα+/KLRG1-) and assigned percentiles from 0 to 100. Individual parts or motif analysis was performed by selecting all CARs that contain a given part of interest. Pairs of parts or motifs analysis was performed by selecting all CARs that contain a given pair of parts. Position analysis was performed by selecting all CARs that contain a given part at a position of interest. Distributions for selected CARs were constructed using the HistogramDistribution functionality and smoothed by using the PDF (probability distribution function) functionality to calculate the probability from 2.5^th^ percentile to 97.5^th^ percentile in steps of 5. The mean and standard error of the mean for each distribution was calculated by repeating the above processing for each of 10 neural networks (for predicted array screen data) or for experimental replicates (pooled screen data).

## Acknowledgements

We thank S. Levinson for sharing tumor cell lines; I. Webster of Zenysis Technologies for assistance with processing sequencing data; R. Alameda and other members of the Lim lab for helpful discussions; W. Maestas and J. Fraser for critical reading of this manuscript.

## Author Contributions

K.G.D and W.A.L. designed research. S.W., K.G.D and S.B designed the AI research. K.G.D, H.K.B, M.S.S., S.W., and P.B. performed research. K.G.D., S.W., and W.A. contributed new reagents or analytic tools. K.G.D. and S.W. analyzed data. S.B., S.C. and W.A.L contributed research guidance and supervision. K.G.D. and W.A.L. wrote the paper. All authors edited the paper.

## Data and materials availability

Reagents are available from the corresponding author upon reasonable request. Data, pre-processing and machine learning data analysis codes will be released open source pending approval and are currently available upon reasonable request.

## FIGURE LEGENDS

**Figure S1.**
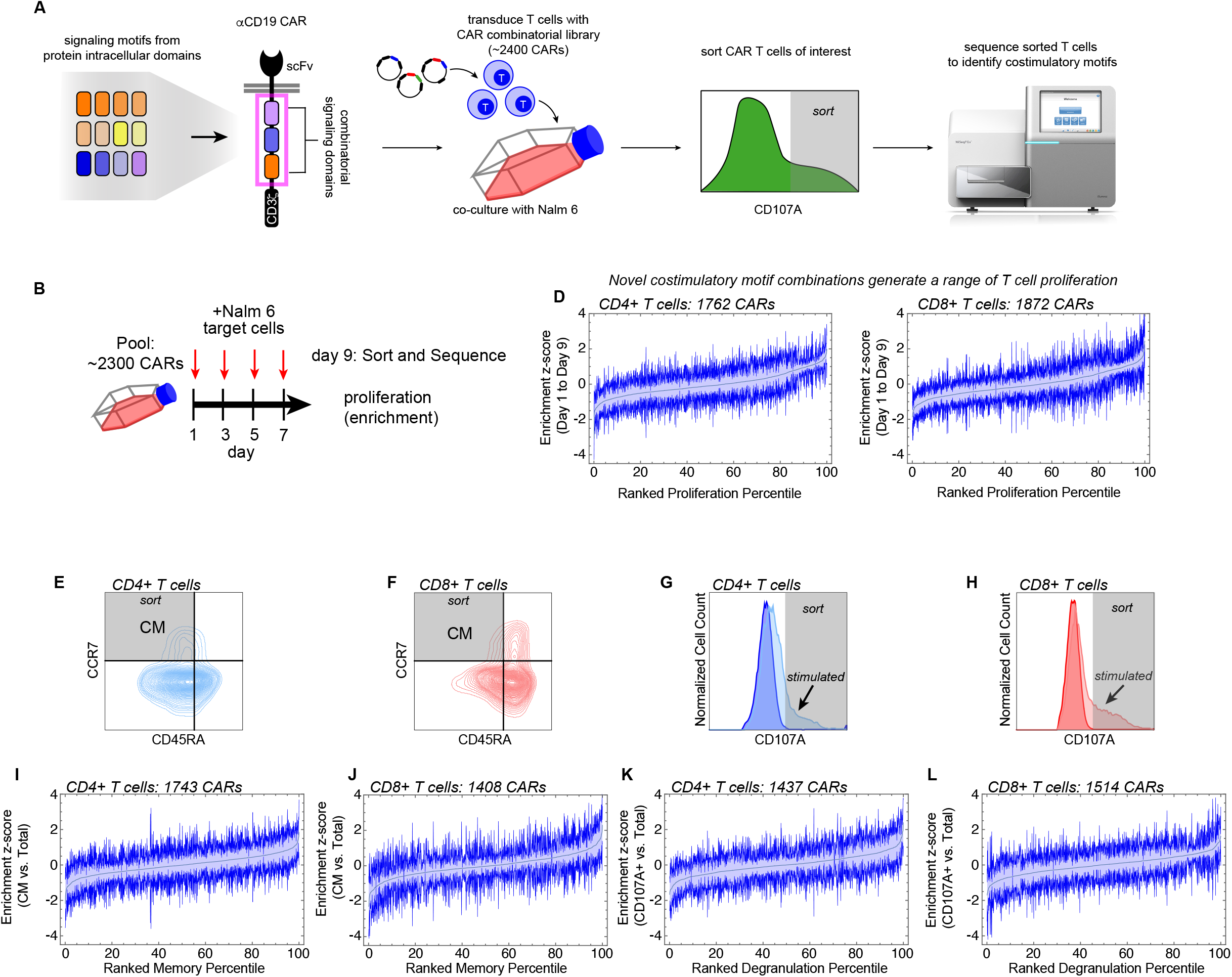
CARs with novel signaling motif combinations generate diverse T cell outputs of proliferation, memory formation, and degranulation in a pooled screen. **A**, Workflow for pooled screening of pooled combinatorial CAR library. **B**, Timeline for pooled combinatorial CAR library screen. **C**, 2378 of the 2379 CAR constructs were detected by sequencing the pooled plasmid library, and over 1700 CAR constructs were detected by sequencing DNA from GFP+ CD4 and CD8 CAR T cells. **D**, CAR T cell proliferation calculated by Log2 fold change in CAR construct frequency 9 days after initial stimulation relative to the starting populations indicates the constructs in the library promote differing degrees of T cell proliferation. **E-H**, Select populations of central memory cells and degranulating cells were isolated by FACS according to the gates shown. Isolated cells were later sequenced. **I-L**, CAR T cell memory formation and degranulation were calculated by Log2 fold change in CAR construct frequency in FACS-isolated select populations on day 9 relative to total populations on day 9. Constructs in the library promote differing degrees of central memory formation and degranulation. All enrichment plot data are mean ± s.e.m. of *n* = 3 replicates.

**Figure S2.**
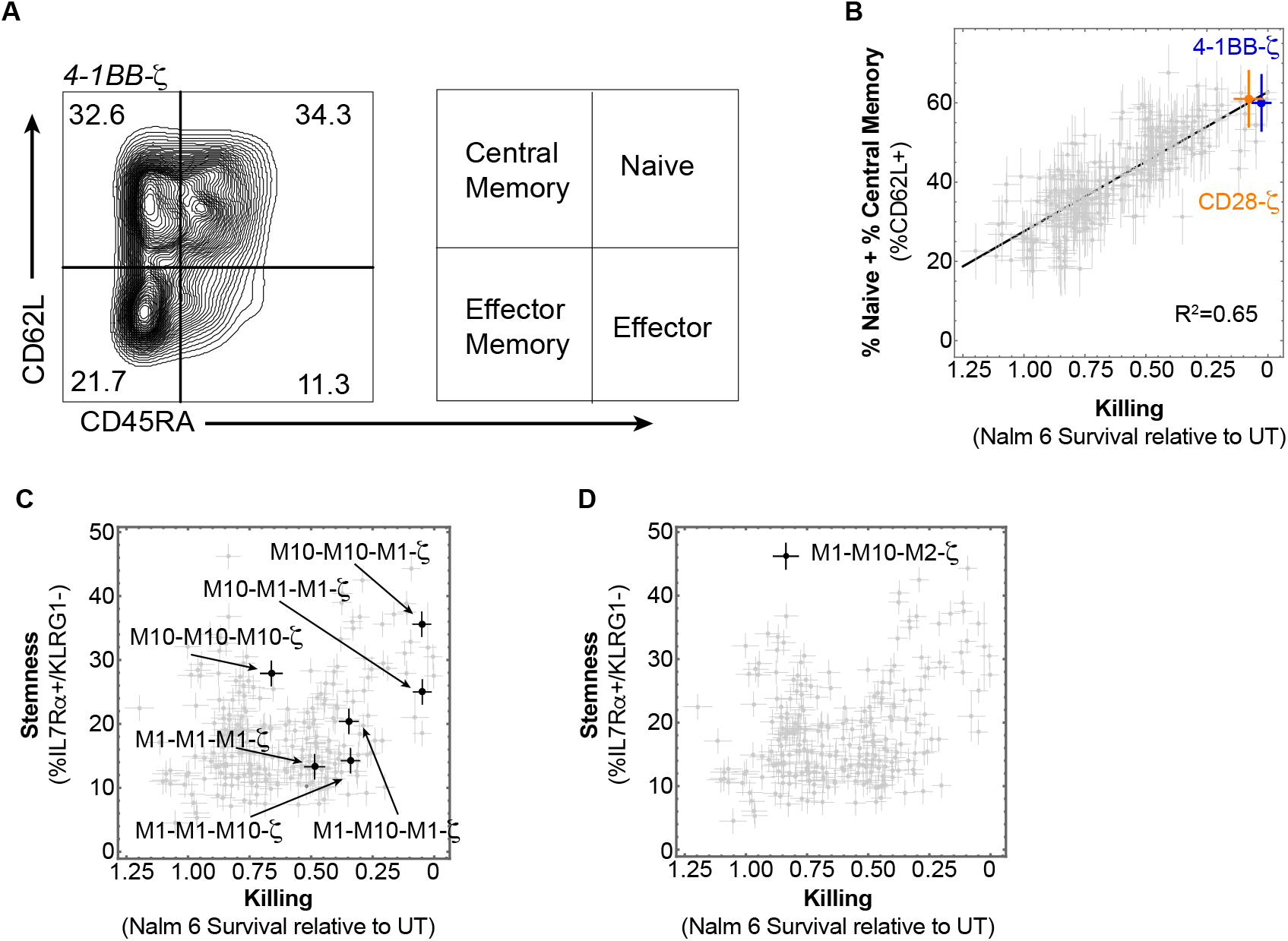
CAR costimulatory domains with novel signaling motif combinations generate diverse cell fates. **A**, CAR T cells with novel signaling motif combinations generate a broad range naïve + central memory populations (quantified by CD62L expression). **B**, CD62L expression is positively correlated with cytotoxicity (r^2^=0.65). Errors for Nalm 6 survival, and %CD62L+ population in **B** were estimated by calculating the average s.e.m. for 7 CAR constructs in the array with internal duplicates. **C**, CARs containing P1 and P10 generate high cytotoxicity and stemness when combined such that P1 is distal from the membrane, but generate reduced cytotoxicity and stemness when not combined or when P1 is not distal from the membrane. **D**. P1-P10-P2-ζ generates low cytotoxicity and high stemness.

**Figure S3.**
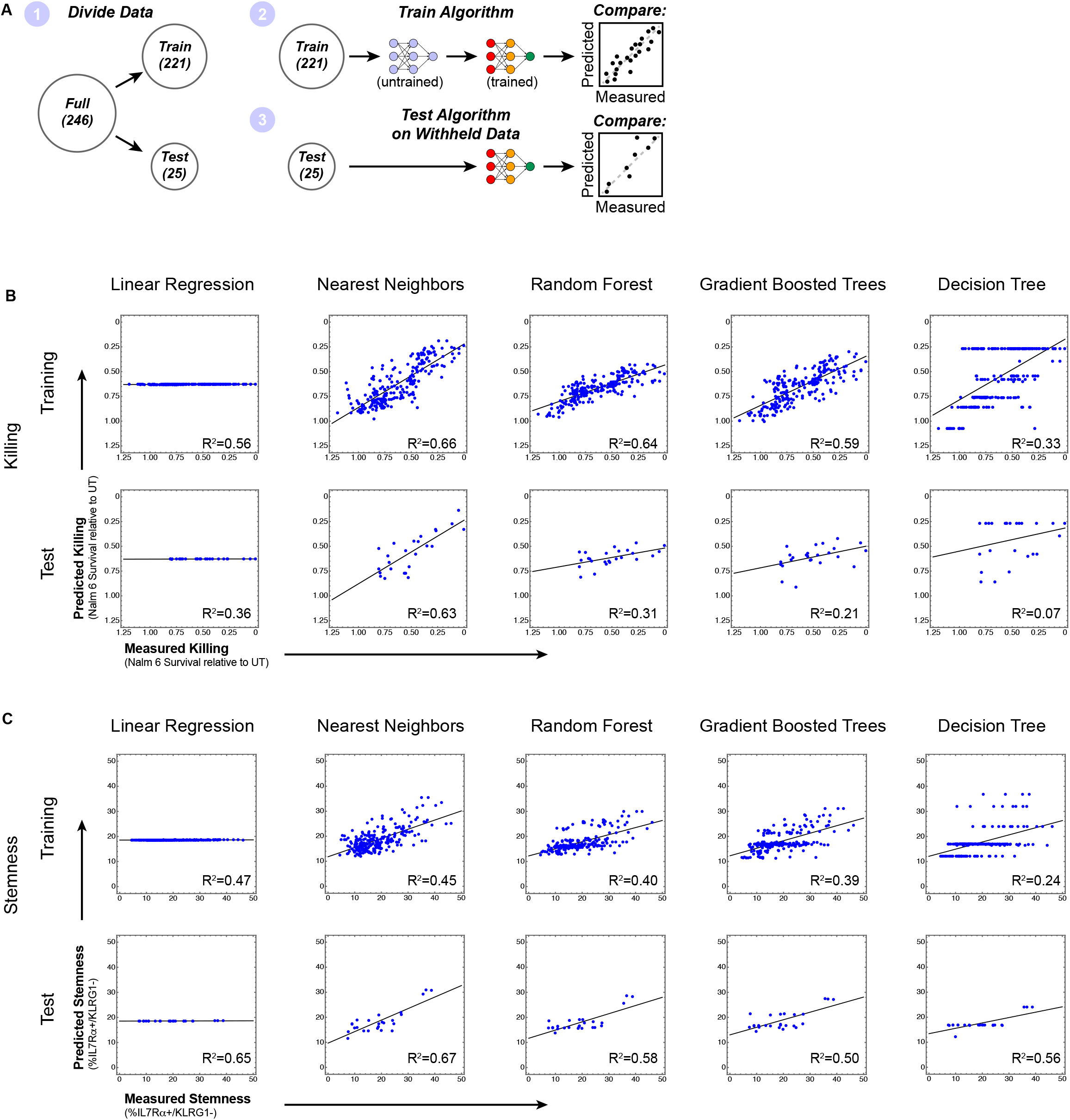
Several common machine learning algorithms fail to predict CAR T cell phenotype. **A**, Array data were subdivided in datasets to train and test various machine learning algorithms. **B-C**, Linear regression, nearest neighbors, random forest, gradient boosted trees, and decision tree algorithms were used to predict cytotoxicity (**B**) and stemness (**C**) resulting from various combinations of library parts.

**Figure S4.**
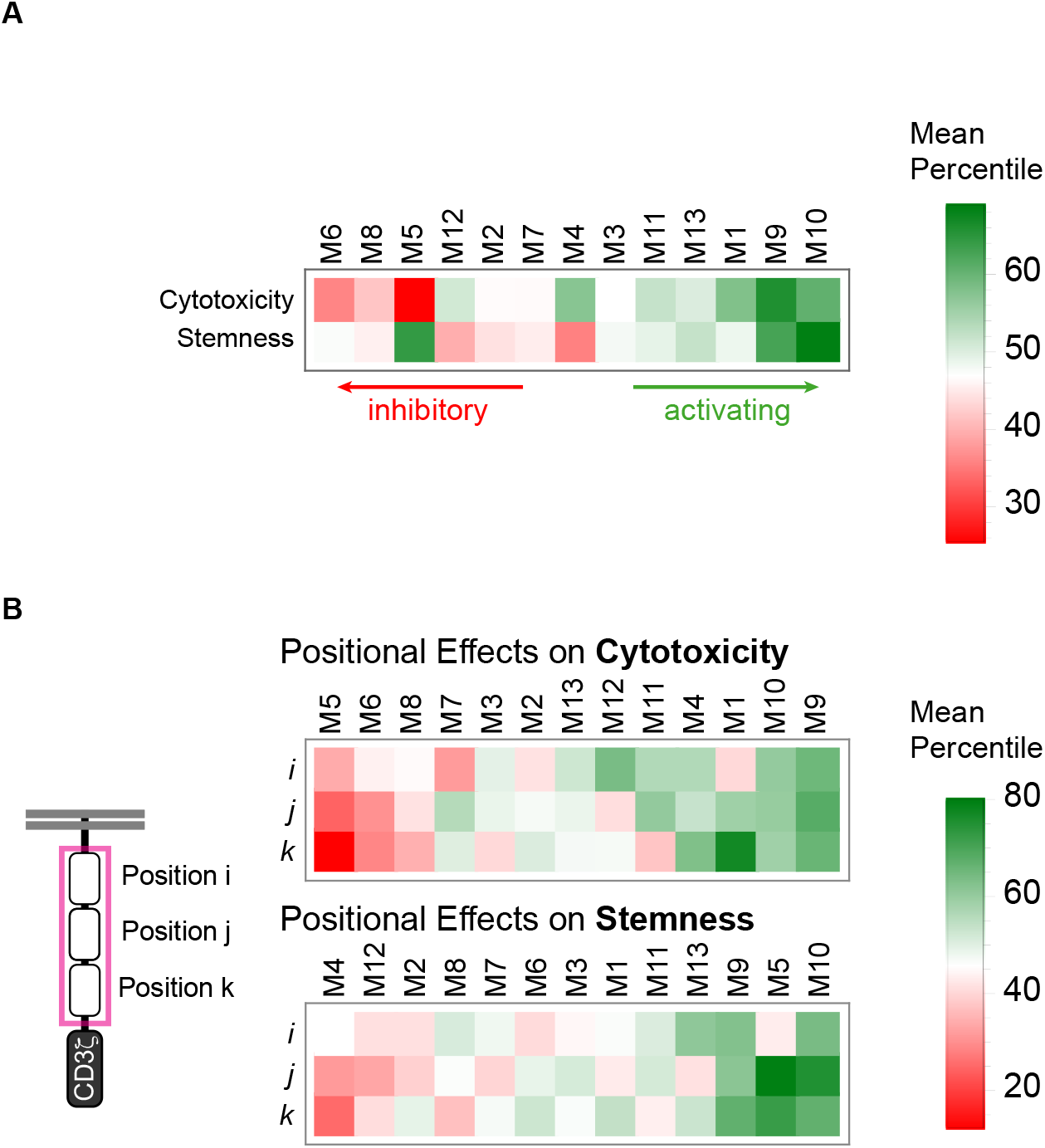
Distribution analysis quantifies elements of linear motif language to extract design parameters for signaling domains. **A**, Heatmaps of mean ranked percentile quantify the overall effects of library parts on CAR T cell cytotoxicity and stemness. **B**, Heatmaps of mean ranked percentile quantify the position-dependent effects of library parts on CAR T cell cytotoxicity and stemness.

**Figure S5.**
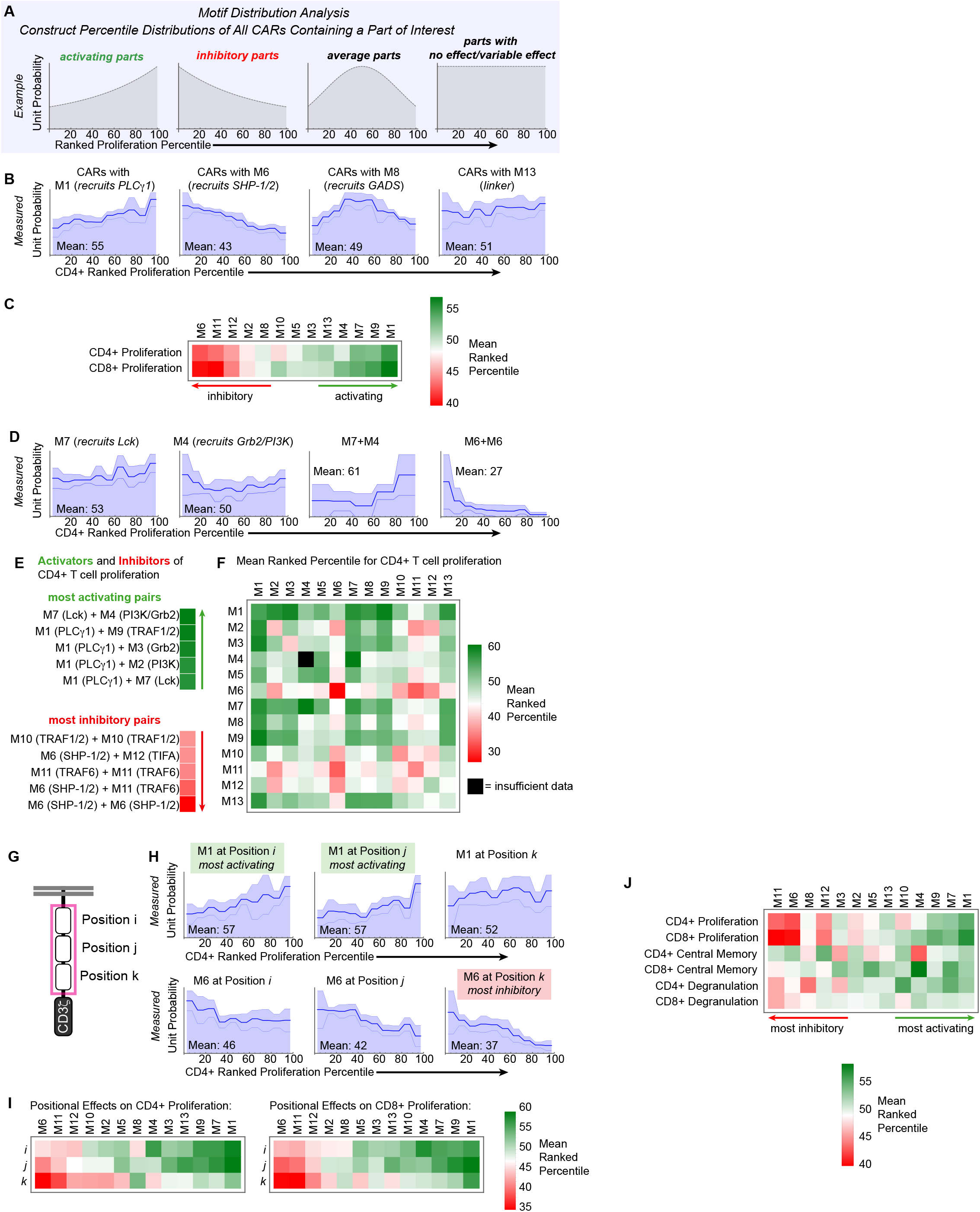
Distribution analysis quantifies contributions of library parts to CAR T cell proliferation in a pooled screen. **A**, Example percentile distributions for CARs that contain parts with various effects on CAR T cell phenotype. **B**, Percentile distributions from pooled screening demonstrate the varied effects of library parts on CD4+ T cell proliferation. The three lines within the distributions represent mean ± s.e.m. for n=3 pooled library screens. **C**, Heatmap of mean ranked percentile quantifies the overall effects of library parts on CAR T cell proliferation measured in pooled screens. **D**, Example percentile distributions for CARs that contain individual parts or pairs parts. **E**, The most activating and most inhibitory pairs calculated using the means of percentile distributions. **F**, Mean ranked percentile for all pairs in the library. **G**, CAR schematic depicting positions *i, j*, and *k* in variable costimulatory domain. **H**, Percentile distributions from pooled screening demonstrate the position-dependent effects of P1 and P12 on CD4+ T cell proliferation. **I**, Heatmaps of mean ranked percentile quantify the position-dependent effects of library parts on CAR T cell proliferation measured in pooled screens. **J**, Heatmap of mean ranked percentile quantifies the overall effects of library parts on CAR T cell proliferation, central memory formation, and degranulation (a proxy for cytotoxicity) measured in pooled screens.

**Figure S6.**
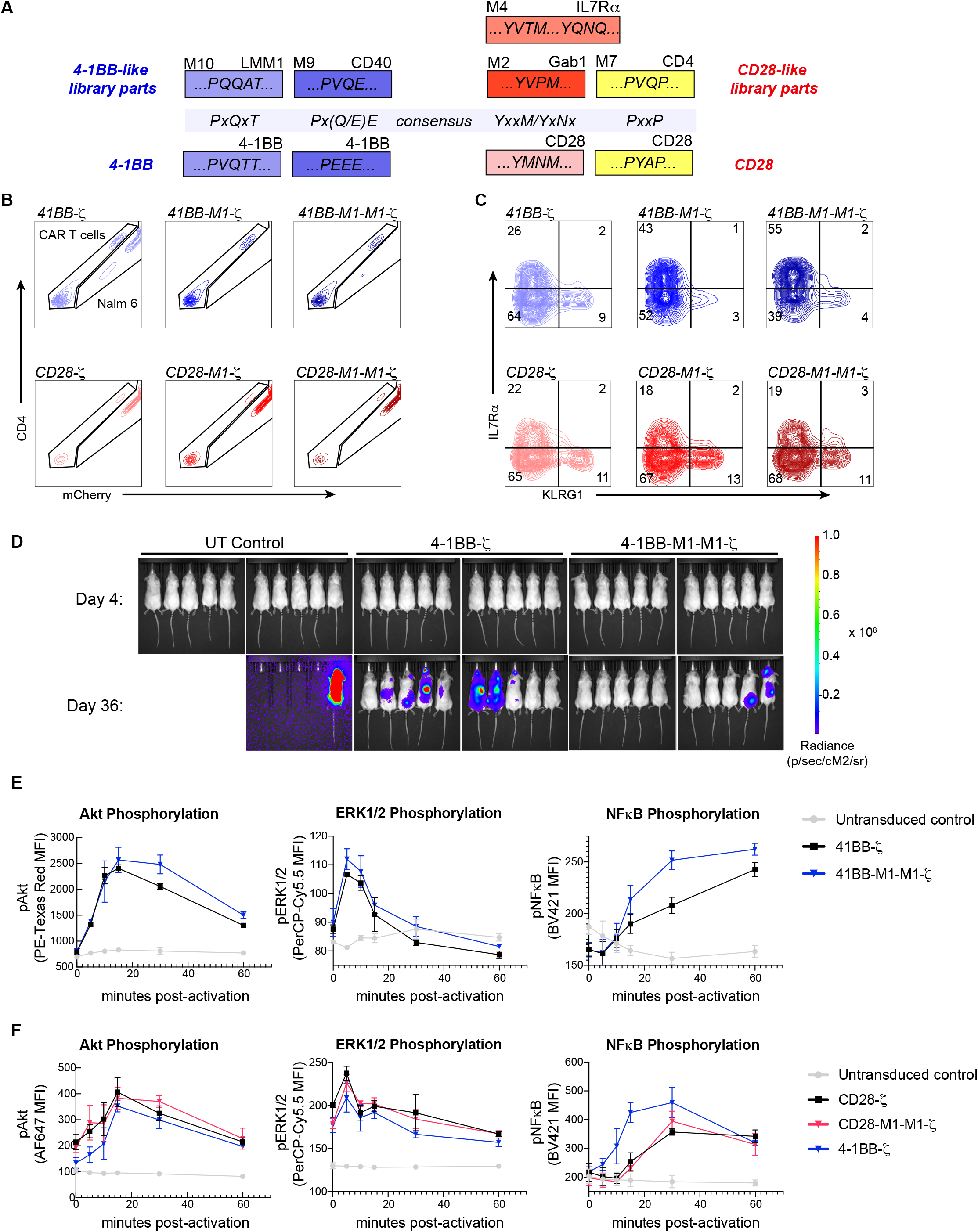
Neural networks accurately predict that PLCγ1 binding motifs improve the cytotoxicity and stemness of 4-1BB-ζ but not CD28-ζ. **A**, Schematics of signaling motifs in 4-1BB, CD28, and functionally similar library parts. **B-C**, *In vitro* assessment of the effect of adding one or two copies of P1 to 4-1BB and CD28 costimulatory domains. Cytotoxicity (B) and IL7Rα and KLRG1 expression (C) were assessed on day 9 after 4 challenges with Nalm 6 target cells. **D**, Tumor progression was monitored using bioluminescent imaging of Nalm 6 expressing the firefly luciferase (fLuc) transgene. Scales are normalized for all time points. **E-F**, Kinetics of phosphorylation upon stimulation of CAR T cells with Nalm 6 target cells, measured by flow cytometry. Kinetic traces represent mean and standard deviation for n=3 replicates.

**Supplemental Table 1.** Hyperparameters for Neural Networks – Cytotoxicity Data

|  |  |  |  |  |  |
| --- | --- | --- | --- | --- | --- |
| Neural Network Model | Convolutional Layer Filters | Convolutional Layer Kernel Size | LSTM Layer Units | Dropout Layer Dropout Rate | Fully Connected Layer Units |
| NN2: | 10 | 2 | 8 | 0.01 | 16 |
| NN3: | 20 | 4 | 8 | 0.2 | 4 |
| NN4: | 10 | 2 | 4 | 0.01 | 64 |
| NN5: | 20 | 4 | 2 | 0.01 | 16 |
| NN6: | 20 | 2 | 4 | 0.01 | 4 |
| NN7: | 10 | 5 | 4 | 0.01 | 16 |
| NN8: | 20 | 5 | 4 | 0.01 | 4 |
| NN9: | 20 | 2 | 4 | 0.01 | 4 |
| NN10: | 50 | 5 | 4 | 0.01 | 4 |

**Supplemental Table 2.** Hyperparameters for Neural Networks – Stemness Data

|  |  |  |  |  |  |
| --- | --- | --- | --- | --- | --- |
| Neural Network Model | Convolutional Layer Filters | Convolutional Layer Kernel Size | LSTM Layer Units | Dropout Layer Dropout Rate | Fully Connected Layer Units |
| NN1: | 20 | 4 | 2 | 0.01 | 64 |
| NN2: | 20 | 4 | 2 | 0.1 | 4 |
| NN3: | 10 | 2 | 8 | 0.01 | 64 |
| NN4: | 10 | 4 | 2 | 0.01 | 64 |
| NN5: | 10 | 5 | 8 | 0.01 | 16 |
| NN6: | 10 | 3 | 8 | 0.1 | 64 |
| NN7: | 10 | 2 | 2 | 0.01 | 16 |
| NN8: | 50 | 4 | 4 | 0.2 | 4 |
| NN9: | 10 | 5 | 8 | 0.01 | 16 |
| NN10: | 20 | 2 | 4 | 0.01 | 4 |

